# A monoallelic immunopeptidome atlas of 10 prevalent canine *DLA-88* alleles

**DOI:** 10.64898/2026.10.06.756904

**Authors:** Jiro Miyamae, Kohei Murakami, Hiroaki Hemmi

## Abstract

Characterization of naturally processed MHC class I (MHC-I)-bound peptides is essential for defining allele-specific peptide-binding features and understanding the functional consequences of MHC-I polymorphism. However, immunopeptidome analyses of dog leukocyte antigen (DLA) class I (DLA-I) molecules remain limited and comprehensive monoallelic ligand datasets for prevalent *DLA-88* alleles are still lacking. Here, we established canine melanoma-derived cell populations expressing individual functional *DLA-88* alleles and generated a monoallelic immunopeptidome atlas of prevalent *DLA-88* alleles. Using liquid chromatography-tandem mass spectrometry, we identified 32,702 naturally processed peptides of 7–15 amino acids across 10 *DLA-88* alleles, substantially expanding the available DLA-88 immunopeptidome landscape. These 10 alleles exhibited canonical peptide length distributions and allele-specific binding motifs that reflected variation in binding-groove pseudo-sequences defined from human MHC-I peptide-contact residues. Ligand sharing between *DLA-88* alleles was associated with binding site distance. Incorporation of the comprehensively characterized DLA-88 ligand datasets in this study increased by more than fourfold the number of published canine DLA-I alleles with ligand-characterized reference alleles predicted to have similar peptide-binding features. We also found that antigen-processing signatures were highly conserved with those reported in humans. As a proof-of-concept application, we selected 54 DLA-88-bound peptides identified in our immunopeptidome analysis as candidate peptides for canine cancer immunotherapy by evaluating the normal tissue expression profiles of their source proteins. This *DLA-88* immunopeptidome atlas provides insights into allele-dependent differences in antigen presentation and represents a valuable resource for developing accurate DLA-88-binding peptide prediction tools and supporting peptide-based immunological studies in dogs.

## Introduction

The efficient induction of antigen-specific cytotoxic T lymphocytes (CTLs) is a key factor in controlling cancer development and viral infection. Major histocompatibility complex class I (MHC-I) molecules are cell-surface glycoproteins that play a primary role in inducing antigen-specific immune responses by presenting antigenic peptides to T cells. A single MHC molecule is estimated to bind and present ∼1,000–10,000 unique peptides to T cells, depending on the alleles [1, 2]. MHC-I genes encoding MHC-I molecules are extremely polymorphic. For instance, in humans, MHC-I genes, referred to as human leukocyte antigen class I (HLA-I) genes, are located on chromosome 6 and comprise six loci: *HLA-A*, *HLA-B*, *HLA-C*, *HLA-E*, *HLA-F*, and *HLA-G*. To date, 17,000 alleles carrying non-synonymous substitutions have been so far published at these six loci across human populations (IMGT-HLA release3.64 (https://www.ebi.ac.uk/ipd/imgt/hla/), accessed on 4/30/2026). Among the six HLA-I loci, the classical loci *HLA-A*, *HLA-B,* and *HLA-C* are highly polymorphic, with more than 5,000 alleles reported for each locus. The extensive polymorphisms of classical HLA-I loci are functionally linked to the efficient induction of CTL-mediated immune responses against a broad range of antigens, as allelic variations in the peptide-binding groove strongly affect the repertoire of peptides presented for CTL recognition [3]. Therefore, classical MHC-I molecules play a central role in the induction and restriction of antigen-specific CTL responses and contribute to interindividual differences in CTL immunity in an allele-dependent manner. Consequently, in addition to information on MHC polymorphisms in the target species, a detailed characterization of the peptides presented by each MHC molecule can provide deeper insights into the diversity of disease susceptibility and vaccine responsiveness of the target species.

Liquid chromatography–tandem mass spectrometry (LC–MS/MS)-based comprehensive identification of MHC-bound peptides, known as “immunopeptidome analysis,” is a powerful approach for the comprehensive identification of naturally processed and presented peptides bound to MHC molecules [4]. In humans, large-scale immunopeptidome analyses of classical HLA-I molecules have identified more than 500,000 unique HLA-I-bound peptides across more than 140 alleles prevalent in diverse populations [1, 2]. In addition, computational tools have been developed to predict HLA-I-binding peptides from antigen protein sequences, and their predictive performance has been substantially improved by training neural network-based algorithms on large-scale immunopeptidome datasets [1, 5, 6]. Advances in computational prediction tools have facilitated the efficient and rational selection of CTL epitopes and accelerated the development of peptide-based vaccines, thereby promoting numerous clinical studies that evaluate their safety and efficacy against cancer and infectious diseases in humans [7, 8]. These tools are expected to support the development of epitope-based vaccines in veterinary medicine. Indeed, in bovine MHC-I, a prediction tool for MHC-I-restricted T cell epitopes has already been developed to support the control of infectious diseases such as *Theileria parva* and bovine herpesvirus. Its predictive accuracy has been improved by incorporating bovine MHC-I-bound peptide datasets generated by immunopeptidome analyses of multiple bovine MHC-I alleles [9].

Several genomic analyses of dog MHC (dog leukocyte antigen, DLA) genes have revealed that three DLA class I (DLA-I) loci, *DLA-88*, *DLA-12/88L,* and *DLA-64*, are located on chromosome 12, and one divergent DLA-I locus, *DLA-79*, is mapped to chromosome 18 [10–12]. To date, at least 215 DLA-I alleles have been reported in the IPD-MHC database (version report 3.16.0.0 (https://www.ebi.ac.uk/ipd/mhc/), accessed on 4/30/2026) and the NCBI database (https://www.ncbi.nlm.nih.gov/, accessed on 4/30/2026) [13, 14]. Of the 173 official DLA-I alleles registered in the IPD-MHC database, 139 are assigned to the *DLA-88* locus, highlighting the remarkable allelic diversity of this locus. In addition, previous analyses of DLA-I genetic polymorphisms have demonstrated that *DLA-88* is the most polymorphic locus among DLA-I loci [11, 14–16]. Given the prominent role of highly polymorphic classical MHC-I molecules in antigen presentation and CTL induction, these observations support the notion that *DLA-88* represents a major classical DLA-I locus involved in CTL responses in dogs. Despite the extensive accumulation of DLA-I polymorphism data, experimentally defined DLA-I ligand repertoires remain limited. Although monoallelic immunopeptidome data have been reported for selected *DLA-88* alleles, comprehensive ligand datasets across prevalent canine *DLA-88* alleles are still lacking [17–20].

Here, we aimed to generate a monoallelic immunopeptidome atlas of 10 prevalent *DLA-88* alleles selected based on our previous large-scale DLA genotyping data [14]. For each of the 10 *DLA-88* alleles, immunopeptidome analysis was performed using an engineered canine melanoma-derived cell population highly expressing a single *DLA-88* allele and DLA-88-bound peptides were comprehensively identified. We then characterized the allele-specific motifs of DLA-88-bound peptides and assessed the association between binding-groove pseudo-sequence similarity and ligand sharing among 10 *DLA-88* alleles. We also evaluated natural antigen-processing signatures in dogs, primarily reflecting the preferences of proteasomal degradation of cellular proteins and N-terminal trimming of peptides by endoplasmic reticulum aminopeptidase (ERAP). Finally, Finally, to illustrate a potential application of this DLA-88 ligand atlas, we examined source proteins of identified DLA-88-bound peptides that have been highlighted as targets for cancer immunotherapy in humans and assessed their potential as candidate canine cancer antigens using normal tissue expression profiles from the Dog Genome Annotation (DoGA) expression atlas [21].

## 2 Material and Method

### 2.1 Cells and cell culture

The canine oral melanoma cell line derived from a mongrel, KMeC, was kindly provided by Professor Takayuki Nakagawa (The University of Tokyo) and maintained in RPMI-1640 with L-Glutamine (Nacalai Tesque, Kyoto, Japan) supplemented with 10% FBS, 1× Penicillin-Streptomycin Solution (FUJIFILM Wako Pure Chemical Corporation, Osaka, Japan), and 1×MEM non-essential Amino Acid Solution (Nacalai Tesque).

### 2.2 Genomic DNA and total RNA preparation and cDNA synthesis

Genomic DNA was extracted from KMeC using the QIAamp DNA Mini Kit (QIAGEN, Hilden, Germany) according to the manufacturer’s protocol. The total RNA samples used in the present study were extracted from peripheral blood samples in our previous study [11]. The study design and protocol were approved by the Ethics Committee of the Animal Medical Center of Nihon University (#3-027, November 1, 2022). Total RNA was treated with DNase I (Thermo Fisher Scientific, Waltham, MA, USA), and cDNA was subsequently synthesized using an oligo-dT primer and ReverTra Ace reverse transcriptase (TOYOBO Co., Ltd., Osaka, Japan), as described previously [11].

### 2.3 *DLA-88* genotyping of KMeC

*DLA-88* genotyping of KMeC was performed as described in our previous study [14]. Briefly, *DLA-88* was amplified by polymerase chain reaction (PCR) from genomic DNA using *DLA-88* specific primers. The nucleotide sequences of exons 2 and 3 were determined by Sanger sequencing, and allele assignment was performed by comparing the resulting sequences with known *DLA-88* allele sequences using Sequencher software version 5.0.1 (Gene Codes Corporation, Ann Arbor, MI, USA).

### 2.4 Plasmid construction

The pSpCas9(BB)-2A-GFP (PX458) plasmid (#48138) was purchased from Addgene (Watertown, MA, USA). The pEBMulti-Hyg vector was purchased from FUJIFILM Wako. Single-guide RNA (sgRNAs) targeting canine β*2-microglobulin* (β*2M*) were designed using CRISPRdirect (https://crispr.dbcls.jp/). The forward and reverse oligonucleotides corresponding to the sgRNA target sequence were annealed and cloned into the PX458 plasmid (cβ2M-PX458) digested with BbsI (New England Biolabs, Ipswich, MA, USA). The coding sequences (CDS) of canine β*2M* and *DLA-88* were amplified using cDNA samples for which *DLA-88* alleles were determined in our previous study [11]. The amplicons were purified using ExoSAP-IT Express (Thermo Fisher Scientific) according to the manufacturer’s instructions, followed by a second PCR with primers containing the Gly-Ser linker sequences. Canine β*2M*–*DLA-88* single chain dimers (β2M–88–SCDs) were generated by overlap PCR using the purified second-round PCR products as templates (**Figure 1A**). The resulting β2M–88–SCD fragments were digested with SalI and ClaI (New England Biolabs) and cloned into the pEBMulti-Hyg vector digested with the same restriction enzymes. For PCR amplification, each 20 μL reaction mixture contained 25 ng of cDNA or purified PCR amplicons, 0.4 units of KOD FX DNA polymerase (TOYOBO), 1 × PCR buffer, 0.4 mM of each dNTP, and 0.3 μM of each primer. PCR cycling was performed according to the manufacturer’s recommendations. The oligonucleotides and primers used in this study are listed in **Table S1**.

**Figure 1.**
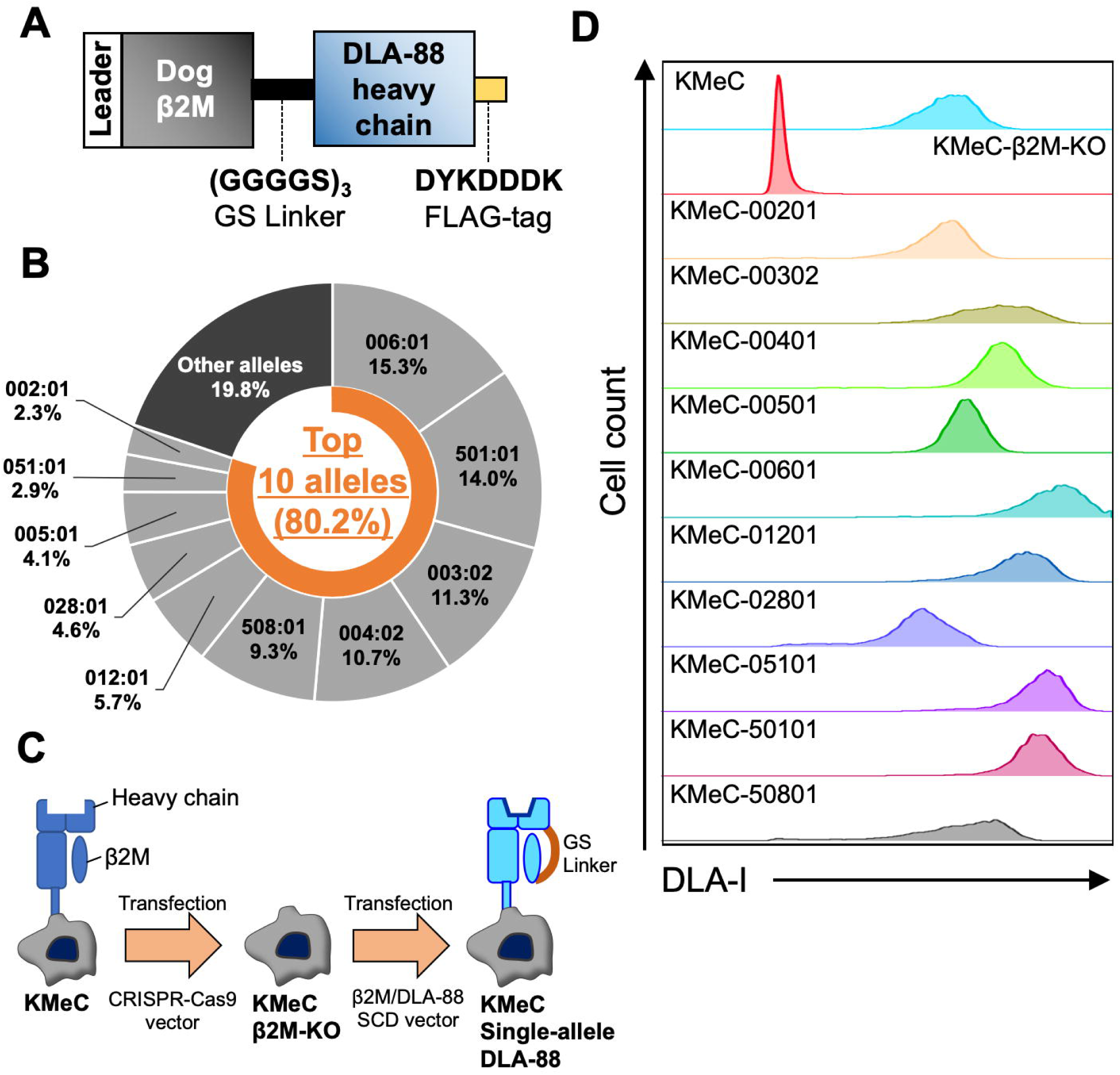
Generation of single-allele *DLA-88*-expressing KMeCs. (A) Schematic representation of the β2M*-*88-SCD construct. (B) Frequencies of the 10 most prevalent *DLA-88* alleles in Japan. The data were summarized from our previous DLA typing study [14] (C) Schematic procedure for generating single-allele *DLA-88*-expressing KMeCs. (D) Flow cytometric analysis of cell-surface DLA-I expression in each single-allele *DLA-88*-expressing KMeC.

### 2.5 Enrichment of single-allele *DLA-88*-expressing KMeCs

To generate KMeC lacking intact cell-surface DLA-I molecules, cβ2M-PX458 was transfected into KMeC using Lipofectamine 2000 (Thermo Fisher Scientific) according to the manufacturer’s instructions. The cell-surface expression of DLA-I molecules was evaluated by flow cytometry using a mouse anti-MHC class I monoclonal antibody (mAb) (clone H58A, Novus Biologicals, Centennial, CO, USA). Cells were stained with the primary mAb at 4°C for 20 min and subsequently stained with PE-conjugated rat anti-mouse IgG2a (BioLegend, San Diego, CA, USA) at 4°C for 10 min. KMeCs lacking cell-surface DLA-I expression were enriched by cell sorting using a FACSAria III cell sorter (BD Biosciences, Franklin Lakes, NJ, USA) and subcloned by limiting dilution. The β2M–88–SCD expression vectors were then transfected into the β2M-dificient KMeC monoclonal cell line using Lipofectamine 2000 reagent, and stable transfectants were selected using 400 μg/mL of hygromycin (FUJIFILM Wako). Cell-surface DLA-I-positive cells were enriched through up to two sequential rounds of cell sorting with a FACSAriaIII cell sorter and maintained in a medium supplemented with 400 μg/mL hygromycin. Enriched DLA-I-positive cells were harvested and washed three times with ice-cold phosphate-buffered saline (PBS). In this study, we refer to these cells as “single-allele *DLA-88*-expressing KMeCs,” indicating that they express a single *DLA-88* allele as a β2M–88–SCD in a β2M-deficient KMeC line background. After counting the cells, the cell pellets were snap-frozen in liquid nitrogen and stored at -80°C until a sufficient number of cells were obtained for subsequent analyses.

### 2.6 Immunoprecipitation of DLA-88/peptide complex

For each experiment, 2–3 × 10^8 single-allele *DLA-88*-expressing KMeCs were used. Cell pellets were gently homogenized by passing through a syringe fitted with an 18G needle repeatedly in lysis buffer containing 50 mM Tris (pH 7.4), 150 mM NaCl, 1 mM EDTA, 1.5% Triton X-100, 1.0% n-Octyl-β-D-glucopyranoside (Dojindo Laboratories, Kumamoto, Japan), 2 mM iodoacetamide (Nacalai Tesque) and a protease inhibitor cocktail (Nacalai Tesque). The lysates were incubated for 1 h at 4°C with rotation and then centrifuged for 10 min at 2,000 g at 4°C to remove large cellular debris. The supernatants were centrifuged for 50 min at 20,000 g at 4°C to remove residual debris and small intracellular organelles. The soluble lysates were filtered through a 0.45 μm filter (Merck Millipore, Burlington, MA, USA) and precleared by incubation with Sepharose 4B beads (Sigma-Aldrich, St. Louis, MO, USA) for 1 h at 4°C with rotation. After centrifugation for 10 min at 2,000 g at 4°C , the cleared lysates were incubated with FLAG M2 affinity gel (Sigma-Aldrich) for 2.5 h at 4°C . The beads were then transferred onto Poly-prep Chromatography Columns (Bio-Rad Laboratories, Hercules, CA, USA) and washed at a flow rate of 1 mL/min at 4°C using the following wash buffers: 5 column volumes (CVs) of lysis buffer, 10 CVs of low-salt wash buffer (20 mM Tris, 150 mM NaCl, 0.1% IGEPAL-CA630 (Sigma-Aldrich)), 10 CVs of high-salt buffer (20 mM Tris, 500 mM NaCl, 0.1% IGEPAL-CA630), 10 CVs of PBS, and 100 CVs of low-salt wash buffer without IGEPAL-CA630. After washing, the DLA-88/peptide complexes were eluted from the FLAG M2 affinity gel with 10% acetic acid. For each DLA allele, *DLA-88*/peptide complex samples were prepared from two to four independent biological replicates.

### 2.7 Isolation, desalting and concentration of DLA-88-bound peptides

*DLA-88*/peptide complexes were heated at 70°C for 10 min to dissociate bound peptides from *DLA-88* molecules. DLA-88-bound peptide fractions were separated from *DLA-88* molecules using Monospin L C18 spin columns (GL Sciences Inc., Tokyo, Japan). Column conditioning, sample loading, and desalting were performed according to the manufacturer’s instructions. DLA-88-bound peptide fractions were eluted with 30% acetonitrile (ACN)/0.1% trifluoroacetic acid (TFA) (Nacalai Tesque). The eluted fractions were dried using a centrifugal evaporator (Tokyo Rikakikai Co., Ltd., Tokyo, Japan). DLA-88-bound peptide fractions and protein fractions were resuspended in 5 μL of 0.1% TFA and 100 μL of low salt buffer without IGEPAL-CA630, respectively, and stored at -80°C .

### 2.8 Determination of peptide sequences with LC-MS/MS

Enriched DLA-88-bound peptide fractions were diluted appropriately and analyzed by nanoLC-MS/MS at the Natural Science Center for Basic Research and Development (N-BARD) at Hiroshima University. Peptide samples were separated chromatographically using Vanquish Neo UHPLC System (Thermo Fisher Scientific) equipped with a NIKKYO NANO HPLC CAPILLARY COLUMN (75 μm inner diameter (i.d.) capillary, 200 mm length, 3 μm C18 particle size; Nikkyo Technos Co., Ltd., Tokyo, Japan) as the analytical column and a PepMap Neo trap column (300 μm i.d. capillary, 5 mm length, 5 μm C18 particle size, Thermo Fisher Scientific). Sample volumes of 1.0−6.6 μL were loaded, and the peptides were eluted at a flow rate of 200 nL/min with a linear gradient of buffer B as follows: 0-4% over 4 min, 4-35% over 96 min, 35-90% over 1 min, and 90% for 4 min. Buffer A consisted of 0.1% formic acid (FA), and buffer B consisted of 0.1% FA in 90% acetonitrile (ACN). The eluted peptides were analyzed using either an LTQ Orbitrap XL (Thermo Fisher Scientific) or an Orbitrap Eclipse Tribrid Mass Spectrometer (Thermo Fisher Scientific). For the LTQ Orbitrap XL, full MS scans were acquired at a resolution of 60,000 for the m/z range of 300–1,500, followed by top 5 data-dependent MS/MS scans using an isolation width of 2.0 m/z and normalized collision energy of 35%. For the Orbitrap Eclipse Tribrid Mass Spectrometer, full MS scans were acquired at a resolution of 30,000 for the m/z range of 380–985. Data-independent MS/MS scans were acquired using sequential 10 m/z quadrupole isolation windows over a precursor m/z range of 380–980. Fragment ions were detected in the Orbitrap over an m/z range of 200–1,800 at a resolution of 30,000, with a normalized collision energy of 28% and a ratio-frequency (RF) lens setting of 30%.

### 2.9 LC-MS/MS data processing

Raw data were analyzed using Proteome Discoverer 3.1 software (Thermo Fisher Scientific) with Sequest HT and INFERYS rescoring workflow [22]. Percolator was used as a validator in this workflow. The INFERYS rescoring node was placed between the Sequest HT and Percolator nodes. The Sequest HT database search was performed against the UniProt database of *Canis lupus familiaris* (TaxID: 9615, Version ID: 2025-02-05), which contains 104,543 protein sequences. The Sequest HT search parameters were set as follows: no enzyme specificity; precursor mass tolerance, 10 ppm; fragment mass error tolerance, 0.8 Da. Oxidation of methionine was set as a variable modification, and carbamidomethylation of cysteine was set as a fixed modification. The false discovery rate (FDR) thresholds in the Percolator node were set to 0.01 for high-confidence and 0.05 for medium-confidence peptide spectrum matches.

### 2.10 Filtering and curation of peptide sequence data

To reduce redundancy caused by putative in vitro proteolysis during sample preparation, peptides nested within longer fragments of ≥13 amino acids and shorter peptides presumed to originate from cleavage of the same ≥13-amino-acid fragment were excluded from the peptide lists generated using Proteome Discoverer 3.1 software (Thermo Fisher Scientific). Because the high expression level of β2M–88–SCDs in KMeC could increase the abundance of peptides derived from degraded β2M–88–SCDs, peptides annotated as *DLA-88* and β2M proteins were also excluded. In addition, potential contaminant peptides derived from non-specifically enriched proteins during immunoprecipitation were identified by comparing well-documented background contaminants listed in the CRAPome database (https://reprint-apms.org) [23]. Candidate contaminants were extracted from the *Homo sapiens* datasets in the CRAPome database using the following filters: Affinity approach, M2 anti-FLAG; Affinity support, agarose; Instrument type, LTQ-Orbitrap.

### 2.11 Computation of binding site distances

A total of 215 DLA class I alleles, including 153 *DLA-88*, 19 *DLA-88L*, 26 *DLA-12*, 9 *DLA-64*, and 8 *DLA-79* alleles, were collected from the IPD-MHC and NCBI databases. In addition, MHC-I alleles for which naturally presented peptides had been identified by LC-MS/MS were collected from previous studies on humans, non-human primates, mice, cattle, swine, and bats [1, 6, 9, 24–34]. In total, 147 alleles with known ligands were collected from species other than dogs. For all collected alleles, pseudo-sequences consisting of 34 key amino acid residues involved in the formation of the MHC peptide-binding groove were generated from their amino acid sequences using Geneious Prime 2019.2.1 (Biomatters Ltd., Auckland, New Zealand), according to the definition established for HLA class I molecules [35].

Pairwise binding site distances between each DLA-I allele and the alleles with known ligands were calculated using the Blosum62 based scoring of pseudo-sequence alignments, as described by Tadros *et al*. [6]. The resulting distance scores ranged from 0 to 1, with higher scores indicating greater dissimilarity between the allele pairs. For each DLA-I allele, the minimum pairwise distance to an allele with known ligands was defined as the binding site distance.

### 2.12 Assessment of antigen-processing signatures

To characterize the antigen-processing preferences for the source proteins of DLA-88-bound peptides, 9-mer DLA-88-bound peptides were compared with randomly generated 9-mer peptides. Random unique 9-mer peptides were generated using an in-house Python program from the same source proteins that yielded 9-mer DLA-88-bound peptides. For each peptide, five flanking amino acid residues upstream and downstream of the peptide sequence were extracted from the corresponding source protein sequences, and the frequency of each residue was calculated at each position. The differences in amino acid frequency between DLA-88-bound and randomly generated peptides were then computed, and their statistical significance was evaluated using the chi-square test. P-values were adjusted using Bonferroni correction. The amino acid frequencies within the 9-mer peptide sequences were also analyzed in the same manner.

### 2.13 Assessment of tissue specific gene expression using the DoGA Genome Atlas

Gene expression data from the DoGA atlas were used to calculate and compare relative gene expression levels across tissues [21]. Normalized read counts (norm.count) from the DoGA Expression Atlas (https://expression-atlas.doggenomeannotation.org/dogaatlas/) were obtained and log2-transformed for each gene after adding a pseudocount of 1 to avoid zeros. Z-scores were calculated based on the log2-transformed expression values, and mean Z-scores were computed for each gene in each tissue. Tissues and genes were clustered based on their mean Z-score profiles using Euclidean distance. In addition, the tissue specificity index (tau) was calculated from the norm.count values, as described previously [36]. To further quantify tissue specificity, the specificity measure (SPM) was calculated from the norm.count values for each gene as the ratio of the squared expression in a focal tissue to the sum of the squared expression across all tissues [37].

## 3. Result

### 3-1. Establishment of single-allele *DLA-88*-expressing KMeCs

To select *DLA-88* alleles for establishing single-allele *DLA-88*-expressing cells, we used our previous large-scale DLA typing dataset to calculate the cumulative coverage of dogs carrying the most frequent *DLA-88* alleles [14]. The 10 most frequent *DLA-88* alleles—*DLA-88*006:01*, *DLA-88*501:0*1, *DLA-88*003:02*, *DLA-88*004:02*, *DLA-88*508:01*, *DLA-88*012:01*, *DLA-88*028:01*, *DLA-88*005:01*, *DLA-88*051:01,* and *DLA-88*002:01*—were selected, and 80.2% of the dogs in the dataset possessed at least one of these alleles (**Figure 1B**). KMeCs endogenously harbor *DLA-88*003:04* and *DLA-88*501:01* and express these intact DLA-I molecules on the cell surface. Therefore, to generate single-allele *DLA-88*-expressing KMeCs, the endogenous β*2M* gene in KMeCs was first disrupted by CRISPR-Cas9 mediated genome editing and the loss of cell-surface DLA-I expression was confirmed in the resulting β2M-dificient (β2M-KO) KMeCs (**Figure 1D**). The cells were then transfected with episomal pEBMulti-Hyg vectors encoding β2M–88–SCDs carrying each of the 10 selected *DLA-88* alleles (**Figure 1C**). Cells with high DLA-I cell-surface expression were isolated by cell sorting, and the enrichment of the DLA-I-high population was confirmed for all 10 *DLA-88* alleles (**Figure 1D**).

### 3-2. Identification of DLA-88-binding peptide motifs for each *DLA-88* allele

For each *DLA-88* allele, DLA-88-bound peptides were isolated and purified from the whole-cell lysates of single-allele *DLA-88*-expressing KMeCs, and their amino acid sequences were identified using LC-MS/MS analysis (see Materials and Methods). To validate our peptide datasets, we first compared the amino acid frequency at each position of *DLA-88*501:01*-bound 9-mer peptides with those of previously reported peptides isolated from human cell lines transduced with the *DLA-88*501:01* heavy chain [17]. The features of the peptide sequences in the present study were strongly (correlation coefficient (r) = 0.963) and significantly correlated (p<10^-20^) with those reported previously (**Figure S1A** and **Table S2I**). Next, we examined whether our peptide dataset contained immunoprecipitation-derived contaminants, as certain proteins are known to be repeatedly and non-specifically enriched by FLAG-tag-based affinity purification [23, 38]. Of the 1,696 *DLA-88*501:01-*bound 9-mer peptides identified in this study, 269 were derived from proteins listed in the CRAPome database (https://reprint-apms.org) [23]. However, the amino acid frequency profiles of these 269 peptides were strongly (r = 0.977) and significantly (p<10^-20^) correlated with the remaining 1,427 peptides derived from proteins not listed in the CRAPome database (**Figure S1B**). Moreover, their binding motifs showed no apparent differences, suggesting that the 269 peptides were genuine *DLA-88*501:01*-bound peptides rather than contaminants (**Figure S1C and S1D)**. These results confirmed that our experimental procedure efficiently enriched DLA-88-bound peptides while minimizing non-specific contaminants. Accordingly, peptides derived from CRAPome-listed proteins were retained for subsequent analyses.

A total of 34,490 peptides were identified across the 10 *DLA-88* alleles, of which 32,702 (94.8%) were 7–15 amino acids in length. For each *DLA-88* allele, 2,414–6,216 peptides were identified, and 87.4–99.4% of these peptides were 7–15 amino acids in length (**Figure 2A**). Analysis of the length distribution of 7–15-mer peptides revealed that 9-mers (n = 17,659) predominated across all 10 *DLA-88* alleles, followed by 10-mers (n = 5,654) and 11-mers (n = 4,593) (**Figure 2B**). The complete list of identified DLA-88-bound 7–15-mer peptides is not included in this preprint and will be made publicly available upon publication of the peer-reviewed article. To characterize the allele-specific peptide-binding preferences, we analyzed the putative binding motifs of 9–13-mer peptides for each *DLA-88* allele (**Figure 2C and S2**). Focusing on 9-mer peptides (**Figure 2C**), dominant anchor residues were identified at positions 2 and 9 for all the alleles. At position 9, hydrophobic residues, including leucine (L), valine (V), isoleucine (I), phenylalanine (F), and methionine (M), were predominant (>92%) in six alleles: *DLA-88*002:01*, *DLA-88*003:02*, *DLA-88*004:02*, *DLA-88*006:01*, *DLA-88*012:01,* and *DLA-88*501:01* (**Figure 2C and Table S2A-S2C, S2E, S2F, and S2I**). *DLA-88*051:01* and *DLA-88*508:01* showed similar motifs at position 9, although the frequency of arginine (R) differed between the two alleles (**Table S2H and S2J)**. In contrast, *DLA-88*005:01* and *DLA-88*028:01* showed distinct motifs at position 9 compared with the other alleles. Position 9 motifs were characterized by relatively conserved preferences for hydrophobic residues, whereas position 2 showed greater allele-dependent variation, with notable motif similarity between *DLA-88*002:01* and *DLA-88*012:01* **(Figure 2C and Table S2A and S2E**) and between *DLA-88*004:02* and *DLA-88*501:01* (**Figure 2C and Table S2C and S2J**). In addition, other dominant or subdominant motifs were observed at positions 1, 3, 5, and 7 in an allele-dependent manner. The dominant anchor residues at positions 2 and C-terminus, together with other subdominant residues, were largely retained as allele-specific motifs in 10-, 11-, 12-, and 13-mer peptides (**Figure S2**). Notably, in *DLA-88*002:01*, *DLA-88*006:01,* and *DLA-88*012:01*, new subdominant anchor residues that were not observed in 9-mer peptide motifs appeared at the second position upstream of the C-terminus in 10–13-mer peptides (**Figure S2**). New subdominant anchor residues also appeared at position 8 in 11-mer peptides for *DLA-88*501:01* and *DLA-88*508:01* (**Figure S2B**).

**Figure 2.**
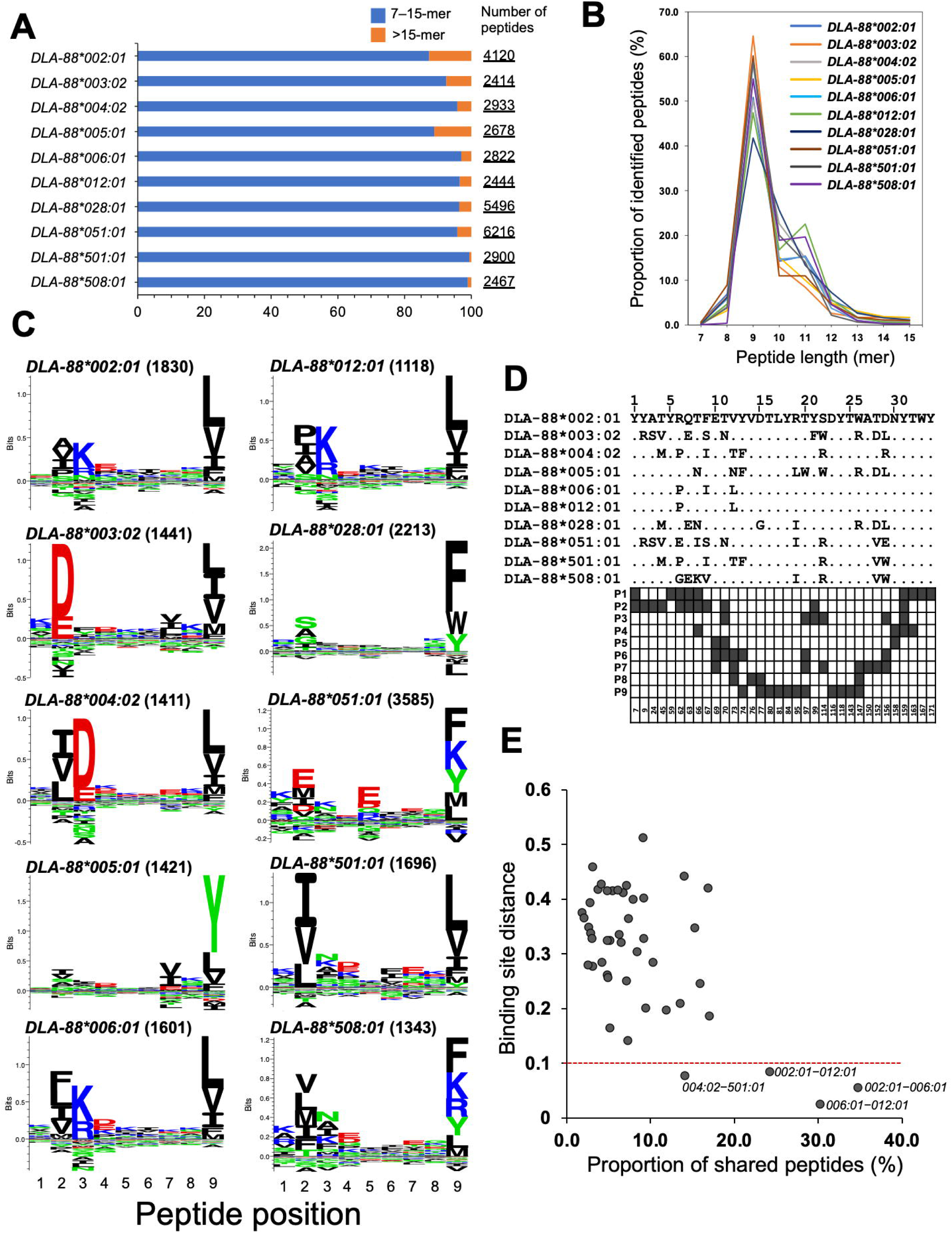
Characteristics of DLA-88-bound peptides identified for 10 prevalent *DLA-88* alleles. (A) Summary of the total number of peptides identified for each *DLA-88* allele. The blue and orange bars indicate peptides of 7–15 amino acids in length and peptides longer than 15 amino acids, respectively. (B) Length distribution of 7–15-mer peptides identified for each *DLA-88* allele. (C) Sequence motifs of DLA-88-bound 9-mer peptides. The number in parentheses following each allele name indicates the number of 9-mer peptides identified for that allele. (D) Alignment of pseudo-sequences of the 10 *DLA-88* alleles. Pseudo-sequences were generated based on 34 key peptide-binding groove residues defined for HLA-I molecules [35]. Dark grey squares in the lower panel indicate the DLA-88 residues predicted to make the closest contact with each position of 9-mer peptides, from P1 to P9. The numbers below the squares indicate the DLA-88 residue positions corresponding to the HLA-I protein numbering system (https://hla.alleles.org/index.html). (E) Relationship between the proportion of shared DLA-88-bound 9-mer peptides and binding site distances among the 10 *DLA-88* alleles. The red dashed line indicates a binding site distance threshold of 0.1. Allele pairs with binding site distances of less than 0.1 are displayed.

### 3-3. Assessment of the correlation between peptide-binding site similarities and DLA-bound peptide motifs

Previous studies have shown that peptide-binding specificities are often shared among MHC-I alleles with similar pseudo-sequences, which are defined by 34 minimal key residues constituting the peptide-binding groove [6, 35]. Accordingly, comparing pseudo-sequence similarity with peptide binding specificity across alleles provides valuable information for generalizing the rules that govern MHC-I-restricted peptide presentations. Therefore, using our large-scale peptide dataset, we assessed whether pseudo-sequence similarity correlated with peptide binding specificity in dogs. We found that the characteristics of the binding motifs were also reflected by the pseudo-sequence similarity among the *DLA-88* alleles. For instance, *DLA-88*004:02* and *DLA-88*501:01* shared identical pseudo-sequence residues at positions predicted to contact positions 2 and 9 of 9-mer peptides, and these two alleles showed highly similar motifs at positions 2 and 9 (**Figure 2C and 2D; Table S2C and S2I**). However, pseudo-sequence residue 29, corresponding to residue 156 in the DLA protein sequence, is located in proximity to position 3 of the 9-mer peptide and differs between the two alleles (Figure 2D), which may explain the observed difference in the motif at position 3.

When assessing the similarity among peptide-binding grooves of MHC alleles, the binding site distance has been recognized as a useful metric. Tadros *et al*. demonstrated, through pairwise comparisons of HLA pseudo-sequences, that MHC-binding peptides can be accurately predicted, even for alleles lacking known ligands, when the binding site distance to an allele with known ligands is less than 0.1 [6]. To examine whether pseudo-sequence similarity is associated with peptide-binding specificity in dogs, we first calculated the binding site distances among the 10 *DLA-88* alleles for which ligands were identified in this study and compared these distances with the proportion of shared 9-mer peptides for each allele pair (**Figure 2E**). Allele pairs with a binding site distance of <0.1 showed substantially higher peptide sharing than those with a binding site distance of ≥0.1:14.1–34.8% (median, 27.2%) versus 2.1– 17.0% (median, 6.2%). These findings suggest that the binding site distance may be useful for inferring peptide-binding specificity among *DLA-88* alleles. Next, we examined the extent to which the *DLA-88* ligand datasets generated in this study expanded the binding-groove similarity coverage of published canine DLA-I alleles. To this end, we generated two reference sets of MHC-I alleles with known ligands. Allele set 1 comprised 149 alleles, including 147 non-canine MHC alleles from various species (see Materials and Methods) and two previously analyzed *DLA-88* alleles (*DLA-88*501:01* and *DLA-88*034:01*) for which detailed ligand datasets suitable for peptide motif and amino acid frequency analyses were available [17, 19]. Allele set 2 comprised 158 alleles, consisting of allele set 1 plus nine *DLA-88* alleles for which ligands were comprehensively characterized in this study. We then calculated the minimum binding site distance between each of the 215 published DLA-I alleles and the alleles in each reference set. When 215 DLA-I alleles were compared with allele set 1, only 13 alleles (6.1%) had a binding site distance of <0.1. In contrast, comparison with allele set 2 increased this number more than fourfold, to 54 alleles (25.1%) (**Figure 3A**). Notably, all 54 alleles that passed the threshold were *DLA-88* alleles. Sequence logos of the pseudo-sequences showed that the nine *DLA-88* alleles whose ligands were comprehensively characterized in this study retained characteristic features of *DLA-88* peptide-binding grooves, whereas *DLA-64* and *DLA-79* alleles displayed distinct pseudo-sequence patterns (**Figure 3B**). Consistent with this, the distributions of binding site distances for *DLA-64* and *DLA-79* alleles were largely unchanged between allele sets 1 and 2, reflecting the dissimilarity of their binding-groove pseudo-sequences to those of MHC-I alleles with known ligands including *DLA-88* alleles. These results indicate that the DLA-88 ligand datasets generated in this study increased the number of canine DLA-I alleles with a closely related ligand-characterized reference allele based on binding-groove pseudo-sequence similarity.

**Figure 3.**
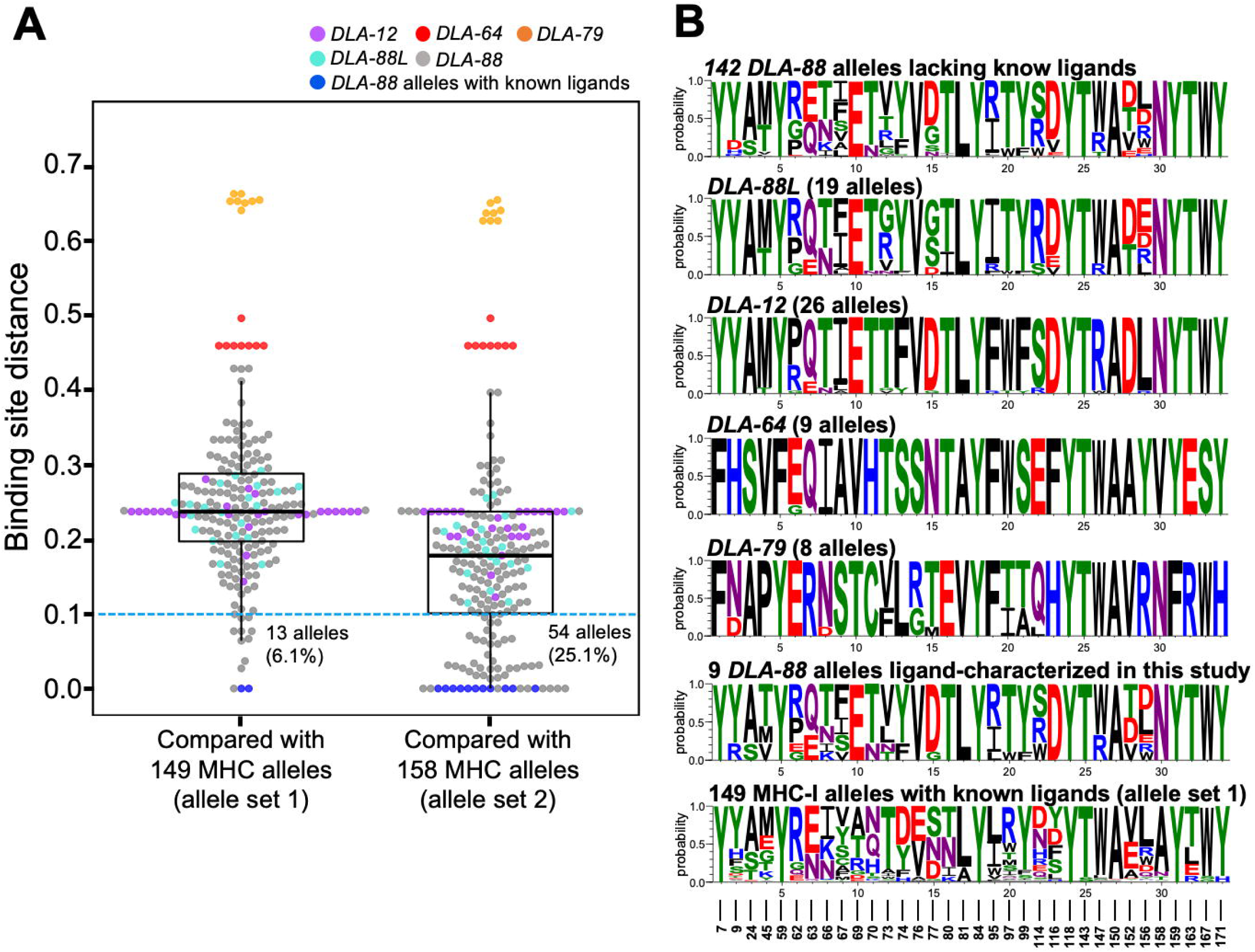
Expansion of binding-groove similarity coverage of canine DLA-I alleles by the DLA-88 ligand atlas. (A) Binding site distance between each canine DLA-I allele and MHC-I alleles with known ligands. Binding site distances were calculated against allele set 1, consisting of 149 ligand-characterized MHC-I alleles including two previously ligand-characterized *DLA-88* alleles (*DLA-88*034:01* and *DLA-88*501:01*) for which detailed ligand datasets suitable for peptide motif and amino acid frequency analyses were available, or alleles set 2, consisting of 158 alleles after addition of nine *DLA-88* alleles whose ligands were comprehensively characterized in this study. Dots indicate individual canine DLA-I alleles and are colored by locus. The blue dashed line indicates a binding site distance threshold of 0.1. The numbers and percentages shown in the graph indicate the number and proportion of DLA-I alleles with binding site distances below 0.1 among all 215 DLA-I alleles, respectively. (B) Sequence logos of the 34-residue binding-groove pseudo-sequences for *DLA-88* alleles lacking know ligands, other canine DLA-I loci, the nine *DLA-88* alleles with ligands repertoires comprehensively characterized in this study, and the 149 MHC-I alleles with known ligand repertoires in allele set 1. The numbers at the bottom indicate residue positions corresponding to the HLA-I protein numbering system (https://hla.alleles.org/index.html).

### 3-4. Characterization of antigen-processing preferences

MHC-I-bound peptides are generated through proteasome-mediated degradation of cellular proteins, followed by N-terminal trimming by ERAPs. The amino acid sequences flanking MHC-bound peptides influence antigen-processing and reflect, at least in part, the substrate specificity of proteasomes and ERAPs [39, 40]. Therefore, we sought to characterize the antigen-processing signatures in dogs using our DLA-88-bound peptide datasets.

In our study, 14,729 unique 9-mer peptides were identified across 10 *DLA-88* alleles. These peptides were derived from 5,839 source proteins with unique master protein accession IDs. To assess the antigen-processing signatures, 15,000 unique 9-mer peptides were randomly generated from the amino acid sequences of the same 5,839 source proteins, and differences in residue frequencies of upstream, intrapeptide, and downstream residues of 9-mer peptides between DLA-88-bound peptides and randomly generated peptides were computed (**Figure 4**). For upstream residues, asparagine (N), alanine (A), and aspartic acid (D) were significantly enriched at the first upstream position (U1) in DLA-88-bound peptides compared to randomly generated peptides, whereas valine (V) was significantly depleted at this position. For downstream residues, alanine (A), glutamine (Q), arginine (R), and lysine (K) were significantly enriched at the first downstream position (D1), whereas acidic residues, including glutamic acid (E) and aspartic acid (D), leucine (L), and tryptophan (W) were significantly depleted. Peptides located at the exact C-termini of the source proteins (labeled with a dash in **Figure 4**) were strongly enriched among DLA-88-bound peptides. A marked depletion of proline (P) was observed at upstream flanking positions U1–U5 and downstream flanking positions D1–D4. This proline depletion extended to the first three residues from both the N- and C-termini of the 9-mer peptides, whereas proline was enriched at the fourth residue from the N-terminus (N4). At the C-terminus of the 9-mer peptides, hydrophobic and aromatic residues, including methionine (M), leucine (L), valine (V), isoleucine (I), tyrosine (Y), phenylalanine (F), and tryptophan (W), were strongly preferred (**Figure 4** and **Table S2**).

**Figure 4.**
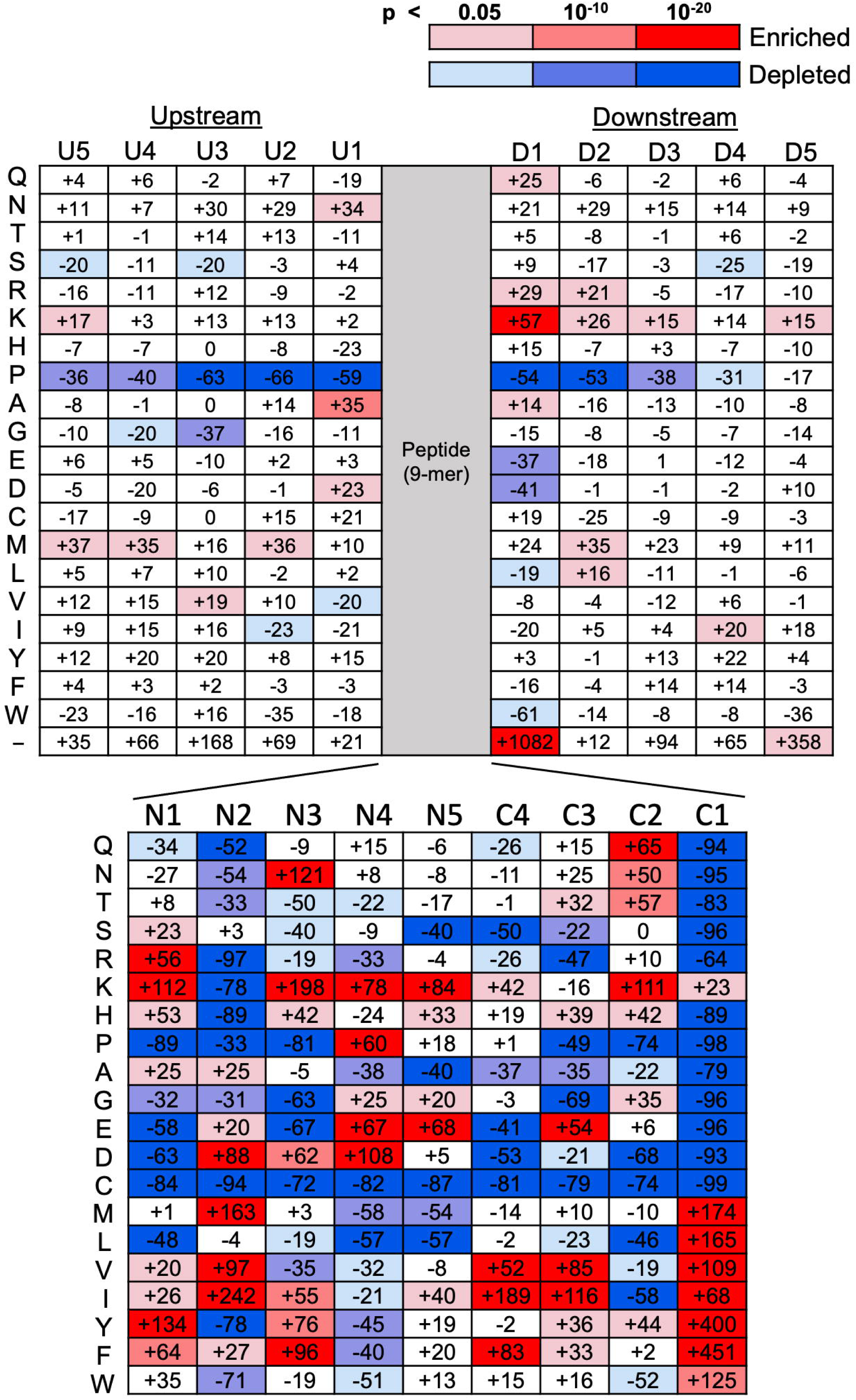
Amino acid enrichment and depletion in DLA-88*-*bound 9-mer peptides and their flanking regions. The heatmap shows the changes in amino acid frequencies in DLA-88-bound 9-mer peptides and their flanking regions compared to those of randomly generated 9-mer peptides derived from the same source proteins. The upper panel shows the percentage change in amino acid frequencies at five upstream (U1– U5) and downstream (D1–D5) flanking positions, whereas the lower panel shows the positions within the 9-mer peptides. N1–N5 and C1–C4 indicate residues at positions 1–5 from the N terminus and positions 1–4 from the C terminus of each 9-mer peptide, respectively. Amino acids are indicated on the left. The dash in the bottom row of the upper panel denotes the absence of a flanking residue at the N- or C-terminus of the source proteins. The numbers within each square exhibit the percentage change in amino acid frequency in the identified DLA-88-bound peptide dataset relative to the randomly generated peptide dataset. Statistical significance was assessed using the chi-square test, and p-values were adjusted using Bonferroni correction.

### 3-5. Screening of peptide candidates for cancer immunotherapy in dogs

From the perspective of comparative oncology, naturally occurring cancers in dogs share many biological, molecular, and therapeutic features with human cancers [41–43]. These similarities suggest that antigens aberrantly expressed and immunogenic in human cancers may also serve as promising candidates for cancer immunotherapy in dogs. As KMeC is a canine melanoma cell line established from primary oral melanoma in a mongrel dog [44], the DLA-88-bound peptide repertoire identified in the present study may include clinically relevant tumor antigen-derived peptides. Therefore, to identify candidate peptides suitable for cancer peptide vaccine development, we screened all detected 9-mer peptides across the 10 *DLA-88* alleles. Fifty-four peptides derived from 18 canine proteins with orthologs overexpressed in various human cancers were identified (**Table 1 and Table S3**) [45–57]. Among the 18 proteins, 10 —AFP, IGF2BP3, SPESP1, ATAD2, CEP55, KIF20A, TPX2, TTK, HSPH1, and MDM2—are listed in **Table 1** because HLA-restricted epitopes derived from these proteins have been reported to elicit T-cell responses or have been evaluated as peptide vaccines in human clinical trials [57–67]. Information on the remaining eight proteins—SPO11, SYCP3, KIF2C, KIF4A, KIF11, KIF14, KIF15, and KIF18A—is provided in **Table S3**. Previous reports have demonstrated the expression of 12 proteins—AFP, IGF2BP3, ATAD2, CEP55, TPX2, TTK, HSPH1, MDM2, KIF2C, KIF11, KIF15, and KIF18A—in various canine cancers or canine cancer cell lines [68–78]. DLA-88-bound peptides shared by multiple alleles were detected for ATAD2, TPX2, MDM2, SYCP3, and KIF11. Two *DLA-88*508:01*-bound peptides, IGF2BP3_343-351_ (IMKKIRESY) and TPX2_244-252_ (KMQQEVVEM), were identical to those previously identified in an immunopeptidome analysis of DH82, a canine histiocytic sarcoma cell line [72].

**Table 1.**
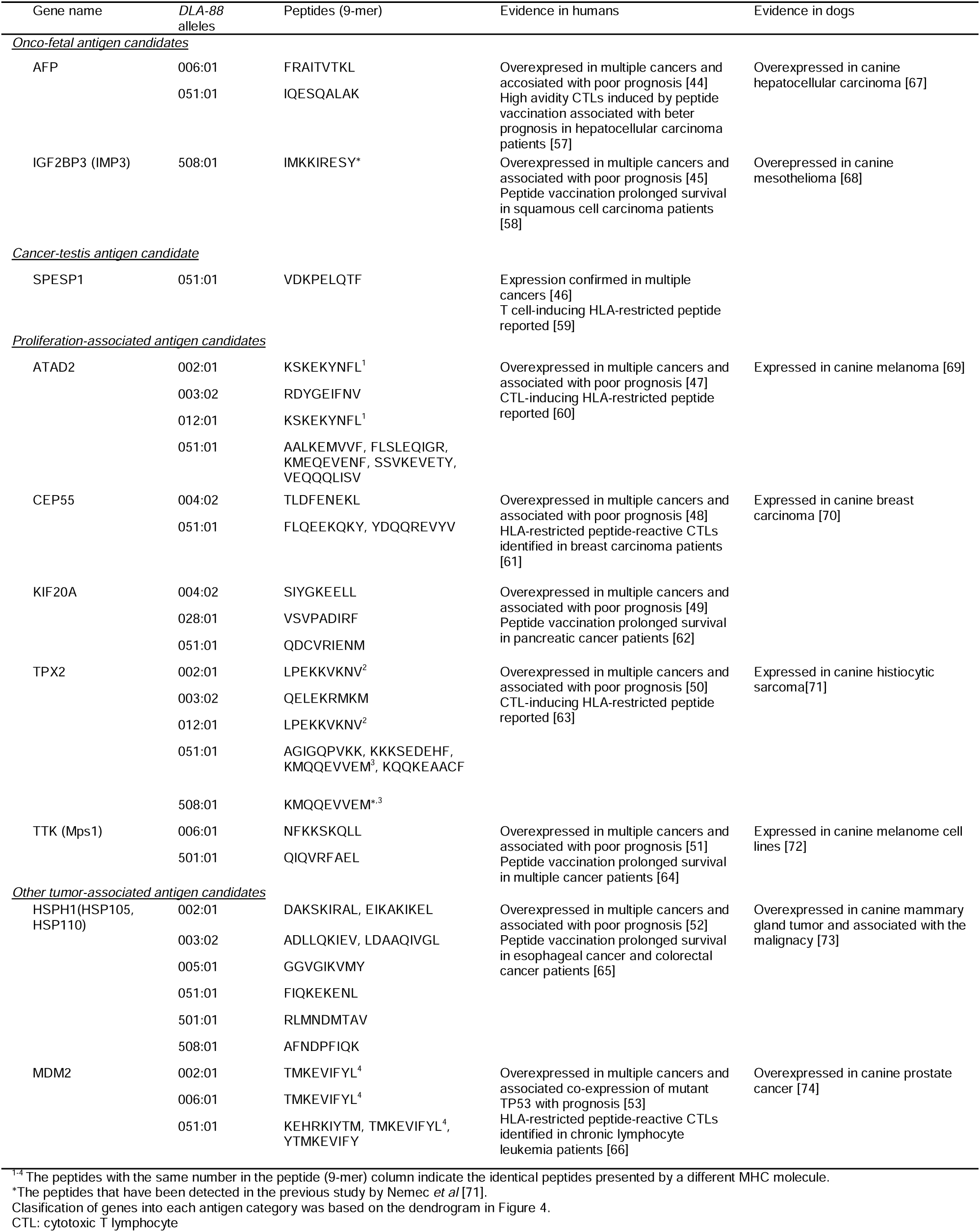
Candidate *DLA-88*-bound 9-mer peptides from human cancer-overexpressed proteins with reported HLA-restricted CTL epitopes.

To evaluate the suitability of these 18 proteins as targets for cancer immunotherapy in dogs, we analyzed their gene expression profiles across 21 representative normal tissues using data from the DoGA Expression Atlas (**Figure 5**) [21], because limited expression in normal tissues is important for minimizing on-target off-tumor toxicity. Hierarchical clustering based on tissue expression patterns classified the 18 genes into four categories: onco-fetal antigen candidates (*AFP* and *IGF2BP3*), cancer-testis antigen candidates (*SYCP3*, *SPESP1,* and *SPO11*), proliferation-associated candidates (*ATAD2*, *CEP55*, *KIF20A*, *TPX2*, *TTK*, *KIF2C*, *KIF4A*, *KIF11*, *KIF14*, *KIF15,* and *KIF18A*), and other tumor-associated antigen (TAA) candidates (*HSPH1* and *MDM2*) (**Figure 5**, **Table 1 and S3**). Although *IGF2BP3* was classified as an onco-fetal antigen candidate, it showed relatively high expression not only in embryonic tissue but also in the lung and retina, resulting in a relatively low SPM value (SPM=0.79). The elevated gene expression of *IGF2BP3* in the lung was consistent with that of a previous study [72]. All 11 proliferation-associated antigen candidates exhibited the highest expression levels in the testes, with comparatively high expression also observed in tissues characterized by active stem cell proliferation, such as the bone marrow and small intestine. *HSPH1* showed the highest expression in the testes, whereas its expression was detected at lower levels in multiple other tissues. *MDM2* was broadly expressed across tissues, similar to *PPIA,* a housekeeping gene. Among the 18 genes, 10—*AFP*, *SYCP3*, *SPESP1*, *SPO11*, *ATAD2*, *CEP55*, *KIF15*, *KIF18A*, *KIF2C,* and *TTK*—exhibited strong tissue-specific expression in dogs, as indicated by high tau and SPM values (tau>0.85, SPM>0.9) [36, 37].

**Figure 5.**
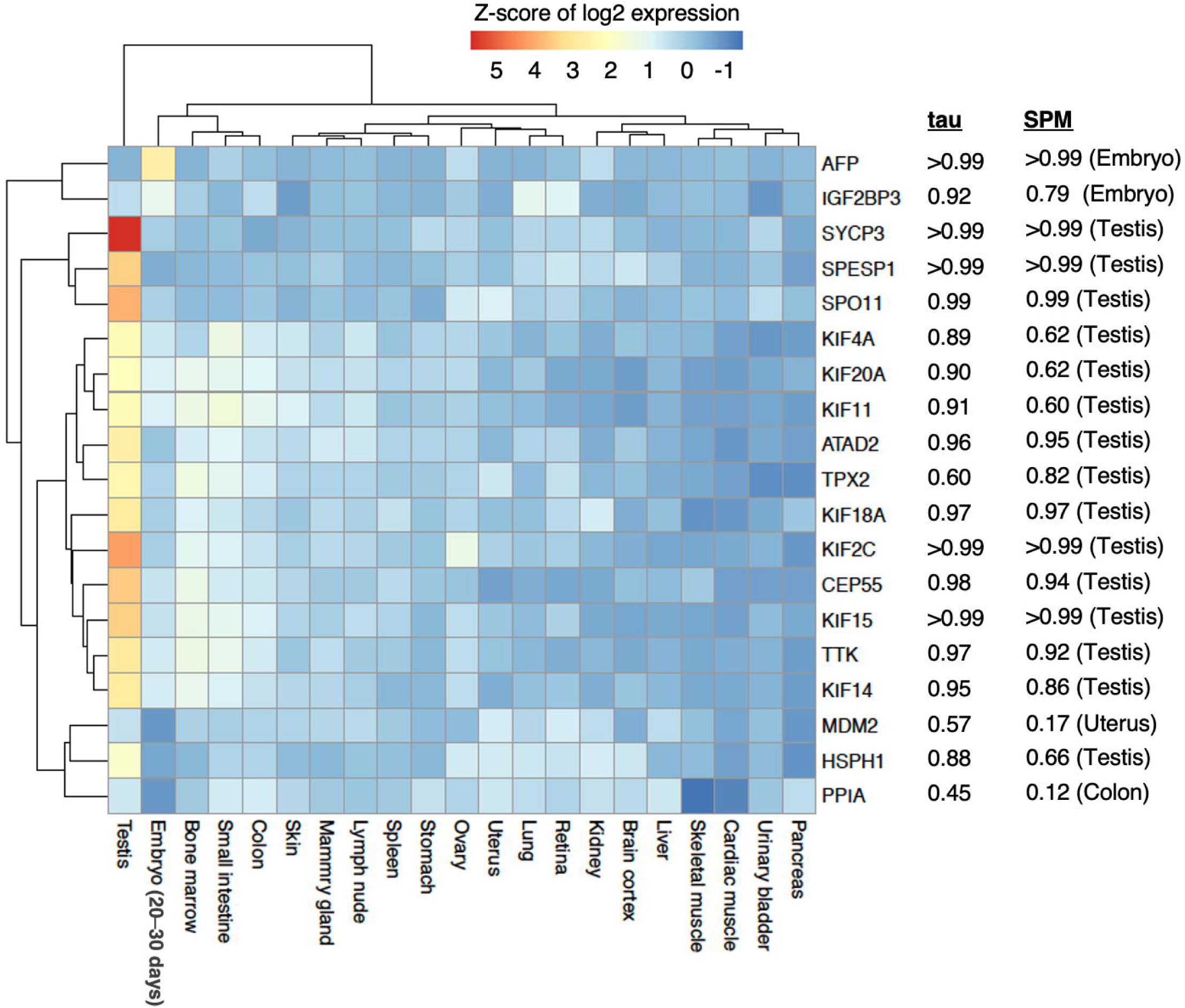
Gene expression profiles of candidate cancer immunotherapy targets across 21 tissues. The gene expression profiles of 18 candidate genes for cancer immunotherapy and the housekeeping gene PPIA were analyzed across 21 representative normal tissues using data from the DoGA Expression Atlas. Tissues are indicated at the bottom of the heatmap, and genes are listed on the right side, together with their tau and SPM values. For SPM, the maximum score for each gene is shown for the corresponding tissue.

## 4. Discussion

In this study, we generated a monoallelic DLA-88 ligand atlas of the 10 prevalent *DLA-88* alleles using engineered canine melanoma-derived cells expressing individual functional *DLA-88* alleles. We identified >2,000 peptides per allele, yielding a total of 32,702 naturally processed peptides of 7–15 amino acids. This dataset substantially expands the currently available DLA-88 immunopeptidome landscape. To our knowledge, this is the first study to systematically compare peptide length distributions, allele-specific motifs, ligand sharing, binding-groove similarity, and antigen-processing signatures across prevalent *DLA-88* alleles using naturally processed and presented ligands.

To date, among veterinary and livestock species other than dogs, immunopeptidome analyses have been reported primarily for bovine and swine MHC-I molecules [9, 34]. These previous studies used cells expressing multiple endogenous MHC molecules and therefore required computational deconvolution for allele assignments. As multi-allelic datasets require motif-based deconvolution, peptides that do not conform to known or predicted motifs may remain unassigned or be excluded during downstream filtering. In contrast, the present study used single-allele *DLA-88*-expressing cells, which enabled the direct identification of peptides bound to individual *DLA-88* alleles and allowed for unbiased characterization of allele-specific binding motifs. In humans, cell lines such as 721.174 and K562, which express cell-surface HLA-I molecules at extremely low levels, are commonly used to generate monoallelic HLA-I-expressing cells for immunopeptidome analyses [1, 28]. However, no cell lines comparable to 721.174 have been established in dogs. Barth *et al*. used human cell lines, K562 and C1R, stably expressing *DLA-88*501:01* and identified more than 3,700 *DLA-88*501:01*-bound peptides [17]. However, because these peptides were identified in the context of human proteins, they may differ from naturally processed peptides in dog cells. Therefore, to obtain peptides naturally processed and presented by *DLA-88* molecules in dog cells, we generated single-allele *DLA-88*-expressing cells by stably expressing β2M–88–SCDs in the canine melanoma-derived cell line KMeC after inactivation of the endogenous β*2M* gene using the CRISPR-Cas9 system. The β2M fused MHC-I SCD is a well-established design that enables a more stable cell-surface MHC-I expression than intact β2M/MHC-I complexes [79]. However, few studies have examined the characteristics of peptides naturally presented by β2M fused MHC-I SCDs. In the present study, the features of peptides bound to β2M–88–SCDs were nearly identical to those of conventional *DLA-88* molecules that were not covalently fused to β2M (**Figure S1A**) [17]. Together, these findings support the utility of β2M fused MHC-I SCD-based single-allele MHC-I-expressing cells for immunopeptidome analyses. Thus, this strategy may be broadly applicable to immunopeptidome analyses in non-model veterinary species for which suitable MHC-low or MHC-deficient cell lines are unavailable, enabling the identification of naturally processed MHC-bound peptides while avoiding the potential biases associated with computational deconvolution.

All 10 *DLA-88* alleles analyzed in this study preferentially bound 9-mer peptides, with 10-mer or 11-mer peptides observed as secondary or tertiary preferences in an allele-dependent manner (**Figure 2B**). Our peptide datasets also included peptides longer than 11-mer from all *DLA-88* alleles (**Figure 2A**), and 12–13-mer peptides largely retained dominant anchor motifs with predominant 9–11-mer peptides (**Figure S2A–S2D**), suggesting that these longer peptides could also be naturally processed and presented on *DLA-88* molecules. The overall length distribution of the DLA-88-bound peptides was consistent with that of classical MHC-I molecules of other species, such as humans, cattle, and swine [9, 28, 34], indicating that all 10 *DLA-88* alleles exhibited a canonical MHC-I peptide length distribution. In HLA-I-restricted antigen presentation, non-canonical long peptides of 13 amino acids or more, derived from viral or cancer antigens, elicit CTL responses. Structural analyses of long-peptide/HLA-I complexes have demonstrated that these long peptides can be accommodated by forming central bulges that protrude from the peptide-binding groove of HLA-I. Such bulged conformations can reshape the peptide/MHC-I surface contacted by T cell receptors (TCRs) and permit atypical patterns of TCR engagement, which, in some cases, may contribute to inducing immunodominant CTL recognition [80, 81]. Therefore, in dogs, as in humans, the exploration of immunogenic peptides should not be limited to canonical 9–11-mer peptides but should also include non-canonical longer peptides. This approach may broaden the discovery of immunodominant CTL epitopes and facilitate the development of peptide-based vaccines for controlling infectious diseases and cancer in dogs.

Recent studies have shown that prediction tools trained on naturally processed MHC ligands achieve higher predictive accuracy than conventional tools trained primarily on binding affinity data obtained from artificially synthesized peptides [1, 5]. Furthermore, incorporating antigen-processing signatures as extrinsic features, in addition to peptide ligand datasets, has also been reported to further improve predictive accuracy [1]. Therefore, the large-scale DLA-88-bound peptide dataset and canine antigen-processing signatures provided in this study are valuable resources for developing highly accurate DLA-88-binding peptide prediction tools. Interestingly, many canine peptide-processing signatures, such as proline depletion at the peptide termini and flanking regions and the enrichment of alanine at U1 and D1 (**Figure 4**), were similar to those observed in humans [1, 28]. This finding suggests that major antigen-processing preferences, likely reflecting proteasomal cleavage and ERAP-mediated peptide trimming, are broadly conserved between dogs and humans.

The binding site distance analysis further supports the utility of the DLA-88 ligand atlas for extending ligand information to alleles lacking experimentally defined repertoires. By comparing published canine DLA-I alleles with MHC-I alleles with known ligand datasets, we found that incorporation of the nine DLA-88 alleles whose ligands were comprehensively characterized in this study increased the number of DLA-I alleles with closely related ligand-characterized reference alleles from 13 to 54 (**Figure 3A**). All alleles that passed this similarity threshold were *DLA-88* alleles, including 43 *DLA-88* alleles without known ligand datasets. These results suggest that our immunopeptidome atlas provides reference alleles for inferring peptide-binding features of a broader set of *DLA-88* alleles. However, binding-site distance alone may not fully capture differences in peptide-binding specificity. As exemplified by *DLA-88*501:01* and *DLA-88*004:02*, peptide-binding motifs can differ substantially even when the pairwise binding site distance is <0.1, particularly when a single amino acid substitution results in a marked change in physicochemical properties, such as polarity and hydrophobicity (**Figure 2C and 2D**). Therefore, future prediction frameworks should incorporate not only pseudo-sequence similarity but also position-specific weights and physicochemical changes informed by DLA immunopeptidome datasets [82]. Such approaches may improve prediction of DLA-88 ligands for alleles lacking experimentally determined ligands.

Although tumor-specific antigens, such as neoantigens, are promising targets for cancer vaccines, most are patient-specific and require personalized approaches. This raises substantial costs and technical challenges, which are likely to limit their broad application in dogs and humans. In contrast, TAAs, including tissue-specific proteins and developmentally associated cancer-germline/testis antigens, are commonly overexpressed in multiple cancer types. These shared TAAs have recently been re-evaluated as promising targets for cost-effective and broadly applicable off-the-shelf cancer vaccines, particularly when combined with immune checkpoint inhibitors [83]. Although TAAs are self-proteins, many clinical studies in humans have demonstrated that TAA-targeting peptide vaccines can induce CTLs capable of recognizing and killing cancer cells without severe toxicity [58, 59, 63, 65, 66]. This is likely attributable to the heightened proliferative state of tumor cells and the inflammatory signatures of the tumor microenvironment. Such aberrant physiological conditions may increase the expression of TAAs and/or antigen-processing machinery, thereby increasing the cell-surface density of TAA-derived peptide/MHC complexes on cancer cells relative to normal autologous cells. In support of this possibility, HLA-I expression has been reported to be higher in cancer cells than in surrounding normal tissues in several early-stage human solid cancers [84]. This enhanced antigen presentation may surpass the threshold required for immune recognition, thereby enabling activated CTLs to efficiently kill cancer cells while minimizing adverse effects [85]. In the present study, 18 canine proteins were identified as candidate targets for cancer immunotherapy from the source proteins of our DLA-88-bound peptides datasets (**Table 1** and **Table S3**). Among these, 17 proteins, excluding MDM2, exhibited the highest gene expression levels in embryonic tissues or testes (**Figure 5**). Of these, 10 proteins—*AFP*, *SYCP3*, *SPESP1*, *SPO11*, *ATAD2*, *CEP55*, *KIF15*, *KIF18A*, *KIF2C,* and *TTK*—exhibited strong embryo/testis-specific gene expression. Moreover, three of these 10 proteins—*AFP*, *KIF15,* and *KIF18A*—have been previously validated as overexpressed in canine neoplastic tissues [68, 77, 78]. These expression profiles may help reduce the risk of adverse effects and promote preferential targeting of cancer cells by CTLs, suggesting that these proteins may be suitable targets for canine cancer immunotherapy. Notably, because *ATAD2* and *TTK* have been reported to be expressed in canine primary melanomas or other melanoma cell lines [70, 73], the 10 peptides derived from these two proteins identified in this study may also be naturally processed and presented by spontaneously occurring canine melanomas (**Table 1**). Further studies are required to demonstrate whether CTLs specific for these peptides can be induced efficiently in tumor-bearing dogs as well as in healthy donors and to evaluate the immunogenicity of these peptides.

Although the 10 *DLA-88* alleles analyzed in this study were selected based on allele frequencies from our previous DLA typing dataset of more than 800 dogs representing 59 breeds in Japan, these alleles do not fully reflect the diversity of the entire dog population [14]. Current information on *DLA-88* allelic diversity is limited, particularly for breeds that are uncommon in Japan. Therefore, further characterization of *DLA-88* allelic diversity, including breeds that have not yet been genotyped, is necessary, and subsequent immunopeptidome analyses focusing on alleles prevalent in the broader dog population, as well as alleles enriched in specific breeds, will be important for understanding the differences in antigen-specific immune responses among dogs.

In conclusion, the present study provides a large dataset of DLA-88-bound peptides for 10 prevalent *DLA-88* alleles, substantially expanding the available DLA-88 immunopeptidome landscape. By systematically characterizing allele-specific peptide-binding motifs, ligand sharing, binding-groove pseudo-sequence similarity, and peptide-processing signatures, this study provides new insights into allele-dependent antigen presentation associated with *DLA-88* polymorphism. Extending this approach to additional *DLA-88* alleles and other DLA-I and DLA-II loci will further improve our understanding of canine MHC immunogenetics. The monoallelic DLA-88 immunopeptidome atlas generated here also provides a foundation for DLA-88 ligand prediction and peptide-based immunological applications in dogs.

## Supporting information

Suppementary Figures

## Acknowledgments

We are grateful to Ms. Tomoko Amimoto of Core Facility Management Center at Hiroshima University for her help in measuring LC-MS/MS data. We thank Professor Takayuki Nakagawa in the University of Tokyo for kindly providing the canine melanoma-derived cell line KMeC.

## Author Contributions

Conceptualization: J.M., K.M., and H.H.; data curation: J.M.; formal analysis: J.M.; funding acquisition: J.M..; investigation, J.M.; methodology, J.M.; project administration, J.M.; resources, J.M.; supervision, J.M., K.M. and H.H.; validation, J.M.; visualization, J.M. writing—original draft, J.M.; writing—review and editing, J.M., K.M. and H.H. All authors have read and agreed to the published version of the manuscript.

## Data availability statement

The mass spectrometry proteomics data have been deposited to the jPOST Repository (JPST004683) and ProteomeXchange Consortium via the jPOST partner repository (PXD079374). These data will be made publicly available upon publication of the peer-reviewed version of this manuscript.

## Declaration of AI or AI-assisted technologies in the writing process

The authors used ChatGPT (GPT-5.5 Thinking, OpenAI) and Paperpal Prime solely to assist with English language editing and to improve the clarity of the manuscript. These tools were not used to develop the scientific content or structure of the manuscript. All AI-assisted outputs were critically reviewed and edited by the authors, who take full responsibility for the final content.

