## Supplementary material for "A monoallelic immunopeptidome atlas of 10 prevalent canine *DLA-88* alleles": Suppementary Figures

**Supplementary figures S1−S2**

**Supplementary figure legends**

**Figure S1. Comparison of amino acid frequencies and peptide motifs for *DLA-88*501:01.***
(A, B) Comparison of amino acid frequencies at each position of 9-mer peptides between the present and previous datasets (A) and between CRAPome-listed and non-CRAPome proteins (B). The correlation coefficient (r) and p-value for the test of no correlation are shown in each graph. The five most frequent amino acids are labeled with their peptide positions in parentheses. (C, D) Sequence logos of 9-mer peptides derived from CRAPome-listed proteins (C) and non-CRAPome proteins (D).

**Figure S2. Sequence motifs of *DLA-88*-bound 10–13-mer peptides**
Sequence motifs are shown for 10-mer (A), 11-mer (B), 12-mer (C), and 13-mer (D) peptides for each *DLA-88* allele. The numbers in parentheses indicate the number of identified peptides used to generate each motif.

### Figure S1

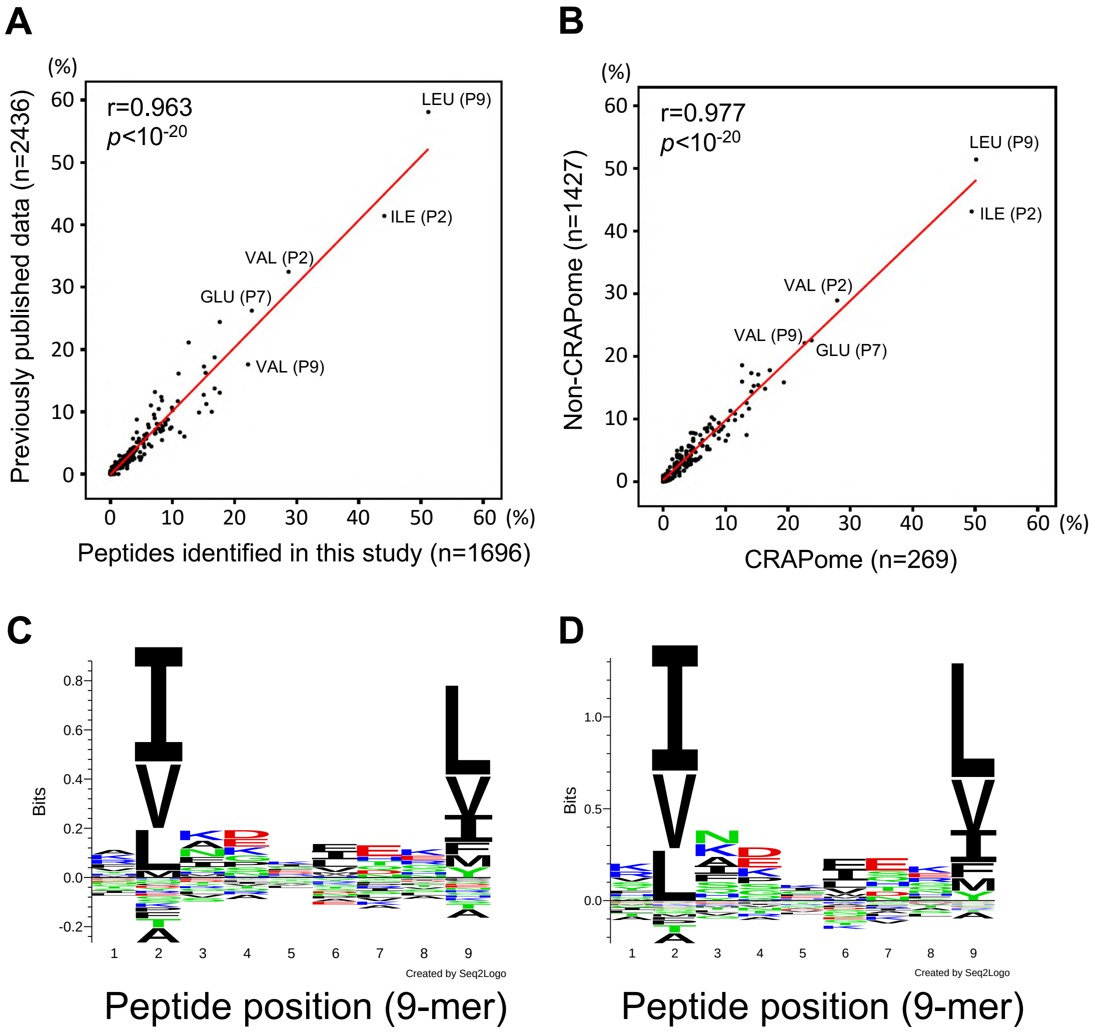
A
(%)
r=0.963
p<10-20
LEU (P9)
ILE (P2)
VAL (P2)
GLU (P7)
VAL (P9)
Previously published data (n=2436)
0 10 20 30 40 50 60 (%)
Peptides identified in this study (n=1696)
B
(%)
r=0.977
p<10-20
LEU (P9)
ILE (P2)
VAL (P2)
VAL (P9)
GLU (P7)
Non-CRAPome (n=1427)
0 10 20 30 40 50 60 (%)
CRAPome (n=269)
C
Bits
1 2 3 4 5 6 7 8 9
Created by Seq2Logo
Peptide position (9-mer)
D
Bits
1 2 3 4 5 6 7 8 9
Created by Seq2Logo
Peptide position (9-mer)

### Figure S2

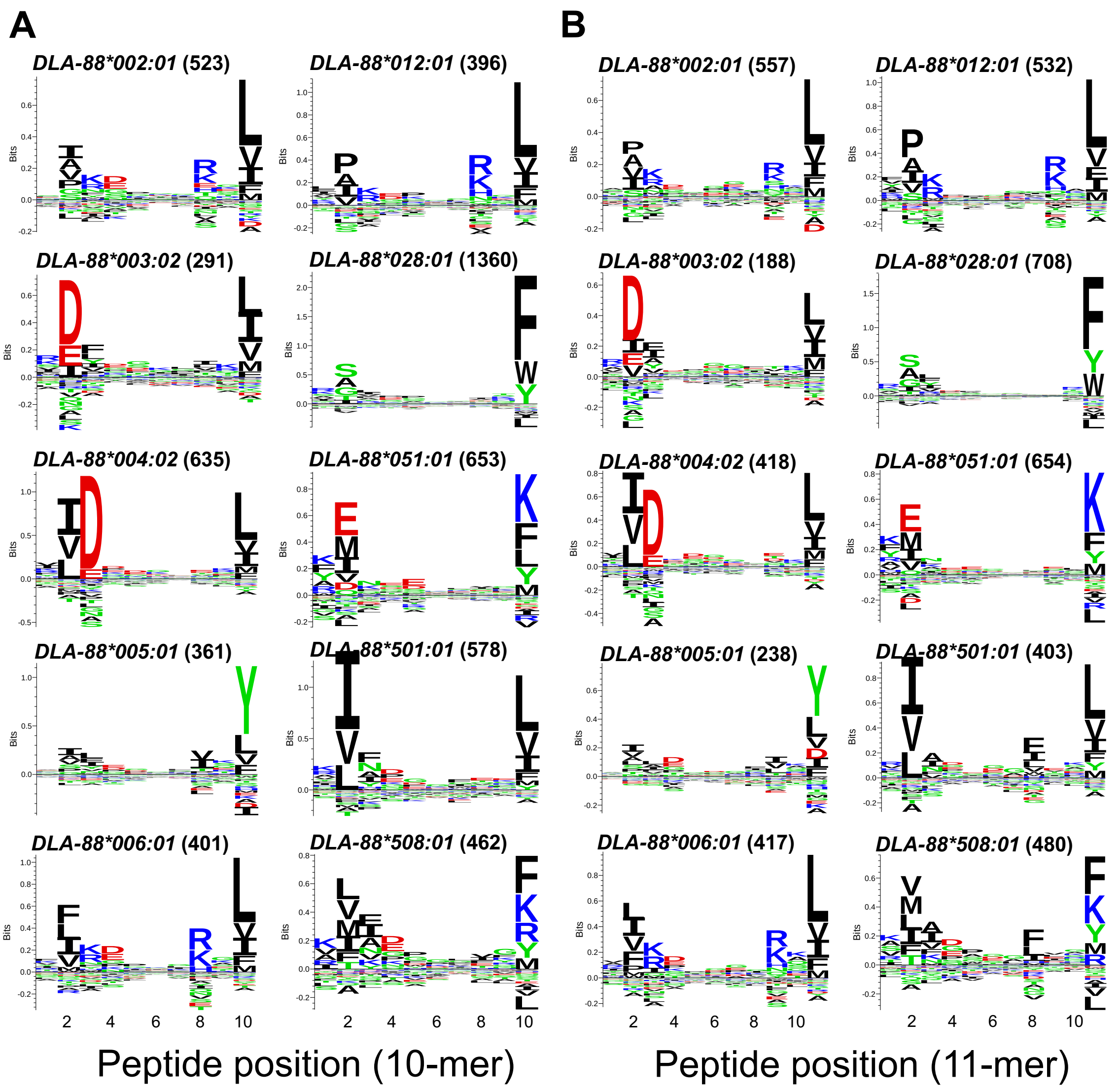
A
B
DLA-88*002:01 (523)
DLA-88*012:01 (396)
DLA-88*003:02 (291)
DLA-88*028:01 (1360)
DLA-88*004:02 (635)
DLA-88*051:01 (653)
DLA-88*005:01 (361)
DLA-88*501:01 (578)
DLA-88*006:01 (401)
DLA-88*508:01 (462)
DLA-88*002:01 (557)
DLA-88*012:01 (532)
DLA-88*003:02 (188)
DLA-88*028:01 (708)
DLA-88*004:02 (418)
DLA-88*051:01 (654)
DLA-88*005:01 (238)
DLA-88*501:01 (403)
DLA-88*006:01 (417)
DLA-88*508:01 (480)
Bits
Peptide position (10-mer)
Peptide position (11-mer)

(continued)

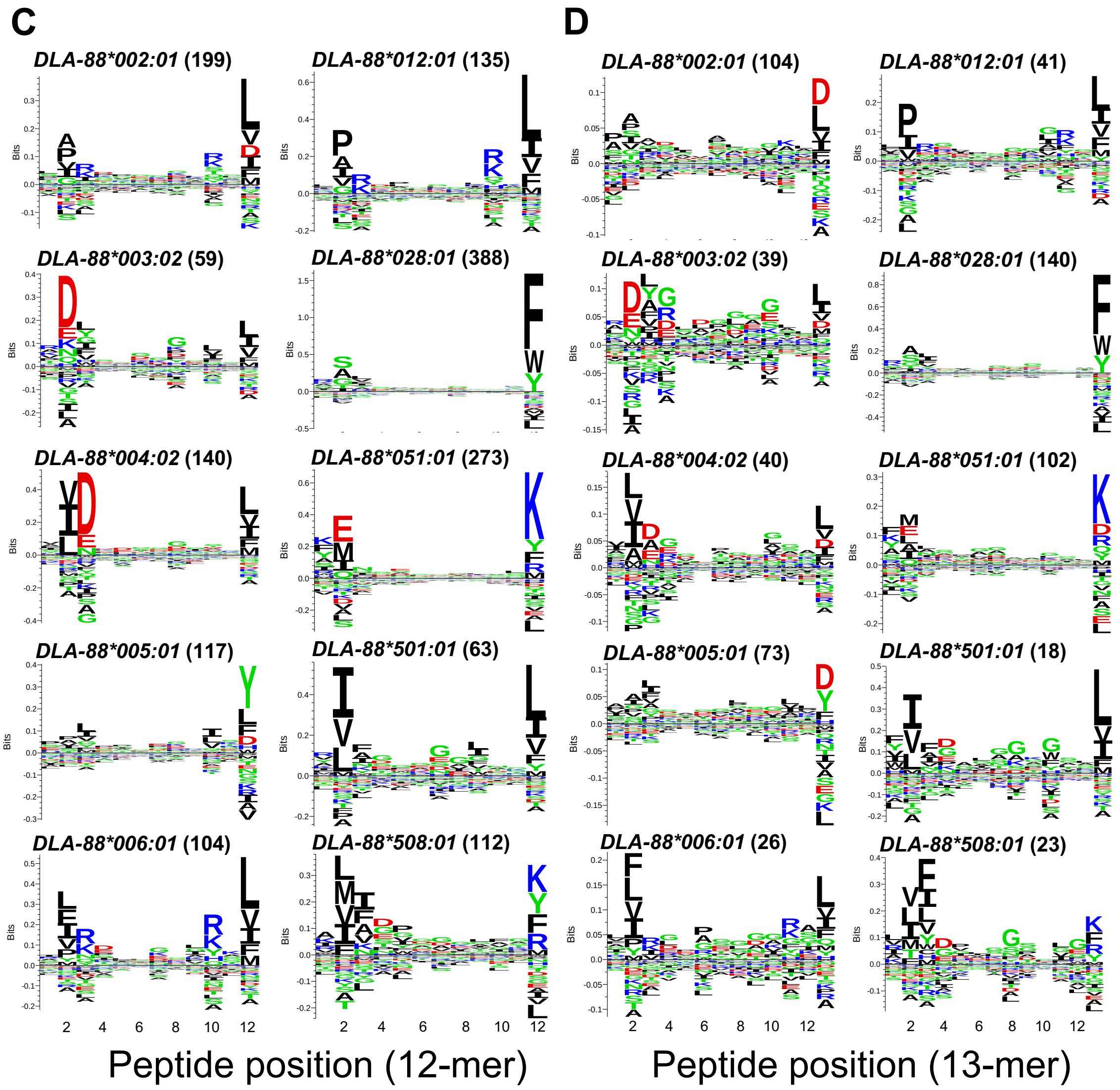
C
D
DLA-88*002:01 (199)
DLA-88*012:01 (135)
DLA-88*002:01 (104)
DLA-88*012:01 (41)
DLA-88*003:02 (59)
DLA-88*028:01 (388)
DLA-88*003:02 (39)
DLA-88*028:01 (140)
DLA-88*004:02 (140)
DLA-88*051:01 (273)
DLA-88*004:02 (40)
DLA-88*051:01 (102)
DLA-88*005:01 (117)
DLA-88*501:01 (63)
DLA-88*005:01 (73)
DLA-88*501:01 (18)
DLA-88*006:01 (104)
DLA-88*508:01 (112)
DLA-88*006:01 (26)
DLA-88*508:01 (23)
Bits
Peptide position (12-mer)
Peptide position (13-mer)
